# Actin Counters Geometry to Guide Plant Cell Division

**DOI:** 10.1101/2025.08.18.670899

**Authors:** Camila Goldy, Samantha Moulin, Yutaro Shimizu, Guillaume Cerutti, Vincent Bayle, Magali Grison, Yvon Jaillais, David Bouchez, Yohann Boutté, Martine Pastuglia, Philippe Andrey, Marie-Cécile Caillaud

**Affiliations:** Laboratoire Reproduction et Développement des Plantes (RDP), Université de Lyon, ENS de Lyon, UCB Lyon 1, CNRS, INRAE, Inria, F-69342 Lyon, France; Laboratoire de Biogenèse Membranaire, Université de Bordeaux, CNRS UMR5200, Villenave d’Ornon, France; Université Paris-Saclay, INRAE, AgroParisTech, Institut Jean-Pierre Bourgin for Plant Sciences (IJPB), 78000 Versailles, France

## Abstract

In multicellular organisms, cell division shapes tissue architecture, cell identity, and function. In walled organisms like plants, division plane orientation irreversibly defines tissue topology and is tightly regulated. While divisions often follow the shortest path enclosing cell volume, certain cells deviate, dividing perpendicular to the growth axis. Tissue-scale mechanical stress has been proposed to guide such orientation, but how these cues are integrated remains unclear. Here, we re-examine the role of the actin cytoskeleton in orienting cell division in *Arabidopsis* root epidermis. Combining cell biology, genetics, pharmacological treatments, 3D segmentation, and modelling, we show that actin is a central molecular actor required to establish cell division orientation against the geometrical rules, highlighting its role in integrating spatial information.

## Main

Accurate positioning of the division plane is essential for proper cell division. It determines whether daughter cells divide symmetrically or asymmetrically and plays a key role in cell differentiation and lineage specification across eukaryotes. In animal cells, division orientation is primarily governed by the mitotic spindle—a dynamic, microtu-bule-based structure that ensures faithful chromosome segregation and guides cleavage furrow placement. Astral microtubules connect the spindle to the cell cortex, interacting with cortical cues and force generators to position the spindle within the cell^1,2^. Concurrently, actin filaments (F-actin) and associated proteins form a supportive cortical scaffold that resists membrane deformation and ensures proper spindle positioning and elongation during metaphase and anaphase^3^. Ultimately, cytokinesis results in the physical separation of the two daughter cells through cleavage. Spindle orientation is tightly regulated by both intrinsic and extrinsic cues (e.g., cell shape, polarity, mechanical forces)^4,5^. Plant cells face unique challenges during division. Surrounded by rigid cell walls, they cannot adopt the rounded morphology typical of mitotic animal cells, and instead rely on plant-specific mechanisms to define the division plane. Most land plant cells lack centrosomes, and the position of the division plane is established before mitosis by the microtubule-based preprophase band (PPB)^6^. Yet, surprisingly, mutants lacking PPBs often exhibit only mild division defects, suggesting the presence of upstream or parallel regulatory mechanisms^7^. Unlike in animal cells, the mitotic spindle plays a limited role in orienting the division plane, and cytokinesis proceeds via formation of a de novo cell plate, guided to a predefined cortical site by the phragmoplast to preserve tissue integrity^8^.

While geometric rules have historically explained plant division orientation—typically aligning the new wall perpendicular to the longest cell axis^9–11^—numerous discrepancies between predicted and observed divisions highlight the limits of geometry alone^12,13^. Mechanical signals, tissue topology, and cytoskeletal organization have emerged as additional layers of control. In particular, cortical microtubules and cell division orientation often align with tensile stress patterns in anisotropic tissues^14^. While the role of the F-actin in orienting plant cell division remains incompletely understood, evidence showed that its disruption alters phragmoplast guidance, tricellular junction positioning, and division plane precision^15–20^, making it a prime candidate to regulate cell division orientation in plants. Here, using the *Arabidopsis* root epidermis as a model, we combine high-resolution microscopy, genetic and pharmacological perturbations, 3D quantitative analysis and modelling to show that actin plays a central role in guiding division orientation, enabling cells to override geometric predictions and integrate positional information during morphogenesis.

### Root epidermal cells do not systematically divide according to area minimisation

The Arabidopsis root meristem is a well-established model for dissecting the molecular mechanisms underlying division orientation^8^. In this system, proliferative divisions occur in a highly stereotypical manner—perpendicular to the root’s growth axis, with new tricellular junctions forming consistently on the lateral faces of the dividing cell^21^, while apicobasal surfaces are rarely used as anchoring sites. However, this tissue is not homogeneous: it comprises distinct, regularly alternating cell types—trichoblasts, which give rise to root hairs, and atrichoblasts, which do not (Fig. 1a,^22^). Despite this predictable patterning, such cellular heterogeneity has largely been overlooked in models of division orientation. To determine whether classical geometric rules—such as divisions forming perpendicular to the growth axis and minimizing new wall area^9–11^—apply uniformly across the epidermis, we performed 3D confocal imaging of calcofluor-stained roots, which labels cellulose-rich walls^21^. Recently-divided cells were identified based on a faint signal at the cell wall between daughter cells, enabling 3D segmentation and volume quantification^21^. Using MorphographX^23^ we calculated the division volume ratio (VR), defined as the volume of the smaller daughter cell relativized to mother cell volume. Most divisions yielded VR values between 0.42 and 0.50 (Fig. 1b, Data S1a), consistent with expectations for near-symmetric divisions^7,24^.

**Fig. 1.**
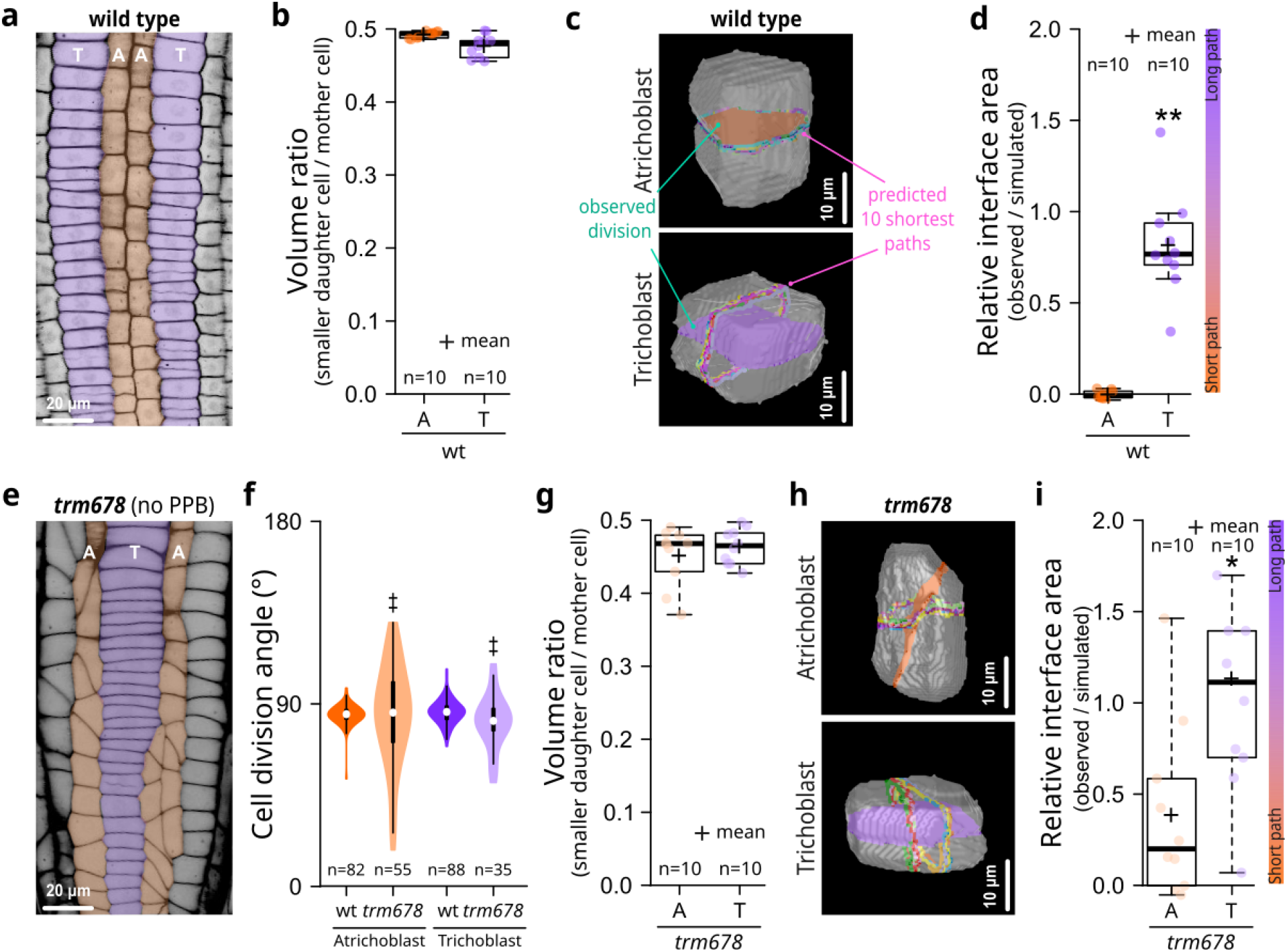
Distinct regulatory mechanisms drive cell division orientation in root epidermal cells. **a.** Representative image of epidermis extracted from a calcofluor-stained Z-stack in WT. Atrichoblast (A, orange) and trichoblast (T, purple) cells. **b.** Quantification of the observed VR in WT. **c.** Representative recently divided, 3D segmented WT epidermal cells. **d.** Representation of the relative interface area between observed and predicted symmetrical divisions in WT. **e.** Representative image extracted from a 3D calcofluor-stained *trm678* root. **f.** Angles between the newly formed cell wall and the main longitudinal axis of the root. **g.** VR quantification in *trm678* epidermal cells. **h.** Representative recently divided, 3D segmented *trm678* epidermal cells. **i.** Representation of the relative interface area between the observed division and the predicted symmetrical cell divisions for *trm678*. * statistical differences; ‡ significant differences in variance distribution.

We tested whether these observed divisions could be predicted by geometry alone. For each reconstructed mother cell, we performed 500 simulations of cell divisions at various VRs by sampling local minima of interface area over all possible division planes^25^ (Fig. 1c). To assess how the observed cell interface area is relative to the simulated local minima interface distribution, we calculate the relative interface area index between the observed area and predicted symmetrical divisions (Fig. 1d). In atrichoblast (A), the observed division planes were oriented following the model’s predicted minimal surface area (relative interface ∽0). This alignment supports the idea that atrichoblasts follow area minimisation when dividing tissues, as previously reported for other, such as the very first cell divisions in the embryo^25–27^. In contrast, trichoblast (T) observed division planes deviated significantly from these predictions, demonstrating that trichoblast divisions do not follow area minimisation (Fig. 1c-d, Data S1a and S2). Instead, trichoblast divide along an alternative path, often perpendicular to the shortest geometric division, a behaviour similar to that recently reported in the underlying cortical layer of the root^21^.

### Directional tissue growth lessens the role of the PPB in guiding cell division orientation

In plants, it is widely accepted that cell division orientation is established before mitosis and is marked by the formation of the PPB, a ring of microtubules that delineates the future cortical division site^28^. However, our findings suggest that trichoblast and atrichoblast cells in the *Arabidopsis* root epidermis may rely on distinct mechanisms to orient their division. To test whether the PPB plays distinct roles depending on cell type, we revisited the phenotype of the *trm678* mutant, which fails to form a PPB due to the loss of three *TON1 RECRUITING MOTIF* genes^7^.

Previous analyses, which did not distinguish between epidermal cell identities, showed that *trm678* roots retained an average division angle close to 90° (relative to the root longitudinal axis), similar to wild-type (WT)^7^. However, the variance around this angle was significantly increased in *trm678*, suggesting reduced robustness in division orientation without the PPB^7^. We examined atrichoblast and trichoblast in longitudinal optical sections to refine these observations. We found that the increased variability in division angle was more pronounced in atrichoblasts than trichoblasts (Fig. 1e). Quantification confirmed that while the mean angle remained near 90° in both WT and *trm678*, the variance was significantly greater in the mutant, especially in atrichoblast cells (Fig. 1f and Data S1b).

By reconstructing recently divided cells in *trm678* using our analysis pipeline, we observed that while daughter cell volume ratios in *trm678* remained close to WT, their variability was slightly increased (Fig. 1g, Data S1a). Strikingly, simulations of division plane orientation in *trm678* revealed that, in atrichoblasts, observed division planes no longer aligned with the shortest geometric path. Instead, divisions appeared to be randomly anchored along the lateral cell surface (Fig. 1h–i, Data S1a and S2). Division orientation in *trm678* trichoblasts remained broadly consistent with WT, though increased dispersion was also observed (Fig. 1h–i, Data S1a and S2). This raised the possibility that both cell types rely less on geometric cues in the mutant background. However, given the differences in cell size and shape between trichoblasts and atrichoblasts, part of the observed variability—particularly in atrichoblasts—may result from physical constraints on division. When normalized by cell length, the effective divergence from idealized division planes may be more comparable between the two cell types, suggesting that both may still implement a similar underlying division strategy, modulated by geometry and cortical anchoring constraints.

### Actin perturbations reset division orientation toward geometry-driven mechanisms

Mutants impaired in PPB formation, such as *TONNEAU 1a* (*ton1a*) and *trm678*, display tilted division planes^7,29^. Nonetheless, division sites in these mutants remain confined to the lateral sides of the cell^7,29^. This observation suggests the presence of additional molecular safeguards that restrict the division plane, preventing it from occurring on the apicobasal axis. Given that (i) disruption of actin dynamics in the shoot apical meristem alters cell geometry, tricellular junction positioning, and division plane fidelity^19^, and (ii) F-actin accumulates at apicobasal cell faces during division in the root meristem^30,31^, we investigated the role of F-actin in orienting divisions in the heterogeneous root epidermis.

Live imaging of the F-actin using plants expressing *LifeAct-tdTom* coupled with the plasma membrane marker *Lti6B-GFP* confirmed a transient accumulation of F-actin at the apicobasal faces of dividing epidermal cells in WT during mitosis (Fig. 2a; Videos S1 and S2;^30^), supporting a potential instructive role for F-actin in orienting the division plane. Perturbation of actin polymerisation using Latrunculin B (LatB) or Cytochalasin D (CytD) treatment significantly reduced the observed apicobasal F-actin enrichment, as quantified via a localisation index (Fig. 2b-c and Data S1c).

**Fig. 2.**
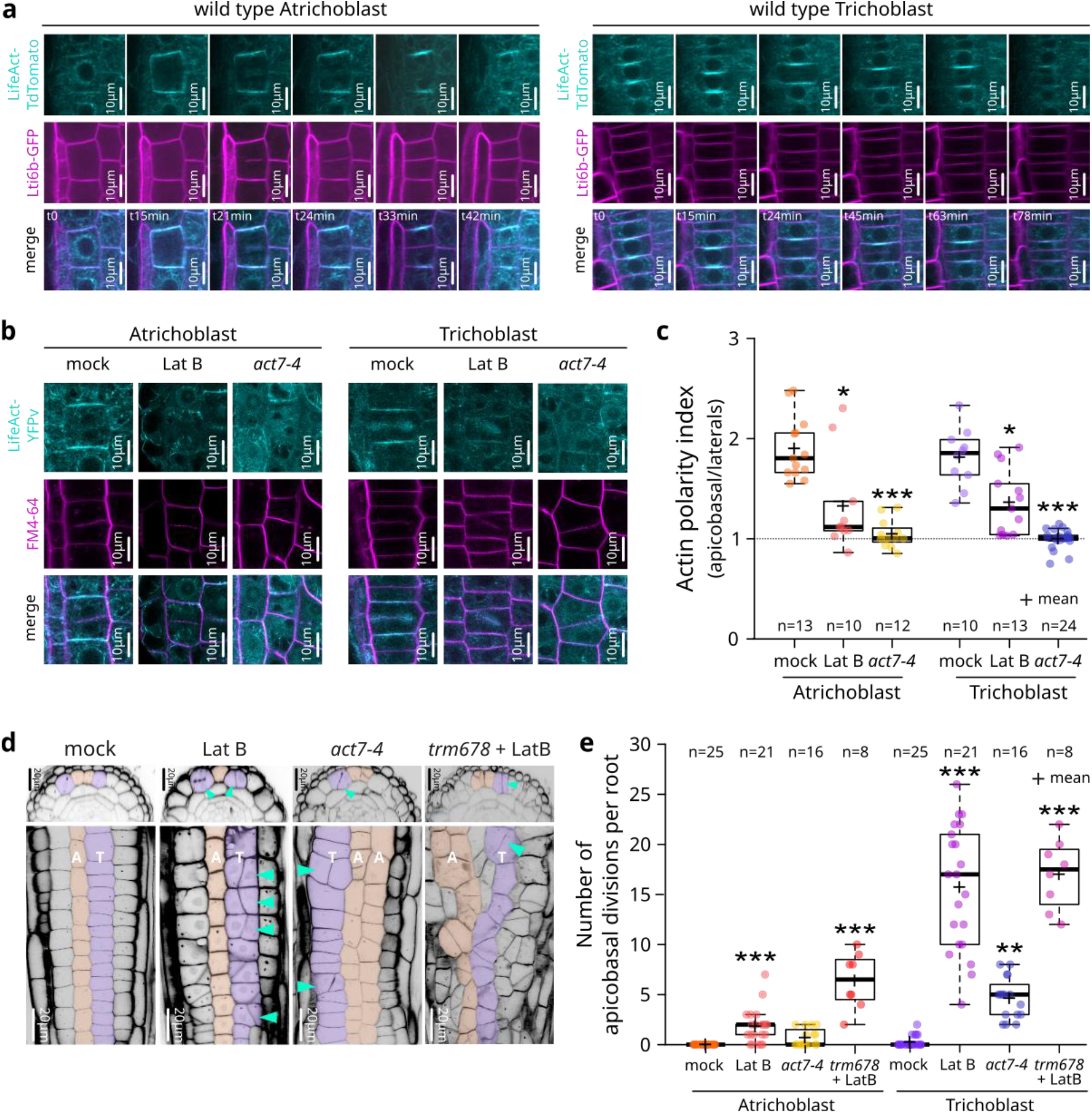
Actin perturbations induce apicobasal cell plate attachment. **a.** Time-lapse of *LifeAct-TdTomato* (cyan) and *Lti6b-GFP* (magenta). **b.** Representative images of LifeAct-YFPv (cyan) and FM4-64 (magenta) in WT 16hmock or -LatB and *act7-4*. **c.** Quantification of localisation index of F-actin between the apicobasal and the lateral faces of dividing epidermal cells in WT 16h-mock or -LatB and *act7-4. N*=3. **d.** Representative image of epidermis layer extracted from a Z-stack of calcofluor-stained WT 16h-mock or -LatB; *act7-4* and *trm678* 16h-LatB. Atrichoblast (A, orange) and trichoblast (T, purple). Turquoise arrowheads point apicobasal divisions. **e.** Quantification of the number of apicobasal divisions per root in WT 16h-mock or -LatB, *act7-4* and *trm678* 16h-LatB. *N*=3. * statistical differences.

Moreover, disruption of F-actin led to striking and reversible defects in division orientation (Fig. 2 and S1). While in WT, mock-treated roots predominantly exhibited transverse anticlinal divisions, actin-disrupted cells frequently divided along abnormal longitudinal anticlinal or periclinal planes (Fig. 2d-e, S1, S2 and Data S1d). Similar apicobasal oriented division defects were also observed in *act7-4* (Fig. 2c-e, S3 and Data S1d), which lack a key actin isoform enriched in dividing root cells^32,33^.

Notably, these defects were significantly more pronounced in trichoblasts—that typically do not divide according to area minimisation—suggesting that the actin cytoskeleton is critical for overriding geometry-based, shortest path division orientations. In contrast, atrichoblasts, which usually follow geometry-based division cues, were less affected by actin perturbation (Fig. 2d-e, Data S1d). Strikingly, LatB treatment of *trm678* roots resulted in a dramatic alteration of meristem architecture. Instead of forming parallel, linear files characteristic of the root meristem, cells adopted a more isotropic topology reminiscent of the shoot apical meristem (Fig. 2d).

By measuring the volume ratio of recent cell divisions in the root meristem we confirmed that the VR observed after division in WT was not affected by the perturbation of the F-actin by LatB treatment or in the *act7-4* (Fig. 3a). Yet, when we simulated cell divisions in both trichoblast and atrichoblast in WT after LatB or in the *act7-4*, all the epidermal cells followed geometrical rule to divide (Fig. 3a and c). Similarly, in *trm678* plants treated with LatB, nearly all cells divide along the shortest path, independent of the organ’s growth axis.

**Fig. 3.**
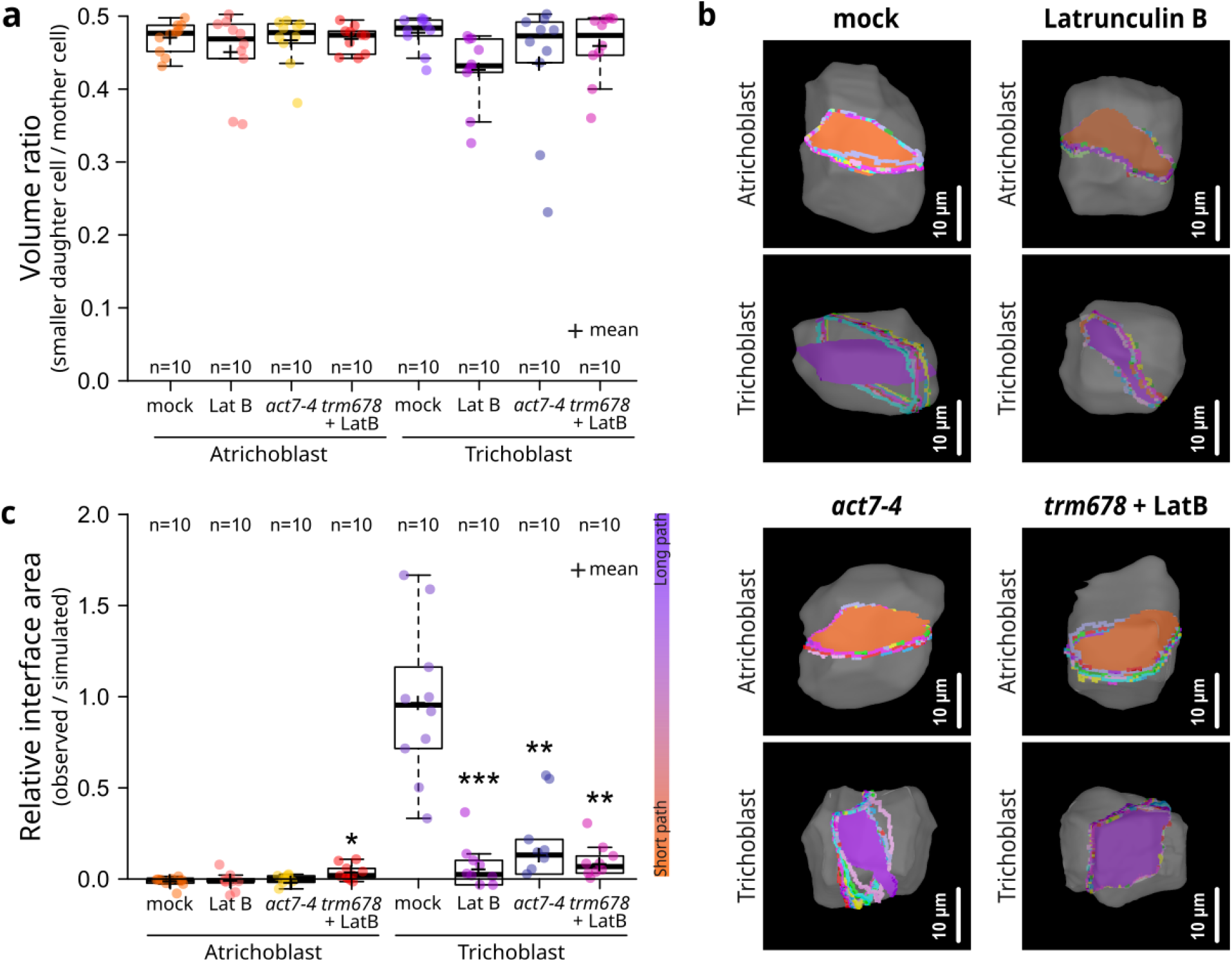
Cells divide along the shortest path when actin is perturbed. **a.** Quantification of the observed VR between the smallest daughter cell and the reconstructed mother cell in WT 16h-mock or -LatB, *act7-4* and *trm678* 16h-LatB. **b.** Representative recently divided, 3D segmented WT 16h-mock or -LatB, *act7-4* and *trm678* 16h-LatB. **c.** Representation of the relative interface area between observed and predicted symmetrical divisions in WT 16h-mock or -LatB, *act7-4* and *trm678* 16h-LatB. Purple and orange planes represent the observed interfaces, dotted colour lines the ten shortest predicted divisions and * statistical differences.

Our findings support a cooperative role between the actin and microtubule cytoskeletons in orienting divisions perpendicular to the growth axis. While the PPB ensures orientation fidelity in specific cell types, actin provides essential spatial cues that reinforce robust patterning, particularly when geometric constraints are not aligned with growth direction.

### Actin perturbation disrupts the alignment between the PPB and the division site

Several proteins, including microtubule-associated factors, are known to accumulate at the future cell division site, where they guide the phragmoplast during cytokinesis and help ensure proper division orientation^8^. To investigate the role of F-actin in maintaining this orientation, we examined how disruption of the actin cytoskeleton affects the localization of these CDZ markers. Following 2h-LatB, the CDZ-localized markers TRM7-3xYPet^7^, YFP-POK1^34^, and IQD8-GFP^35^ consistently remained restricted to the lateral faces of dividing cells, with no detectable signal at the apicobasal surfaces (Fig. 4a–c, Data S1f).

**Fig. 4.**
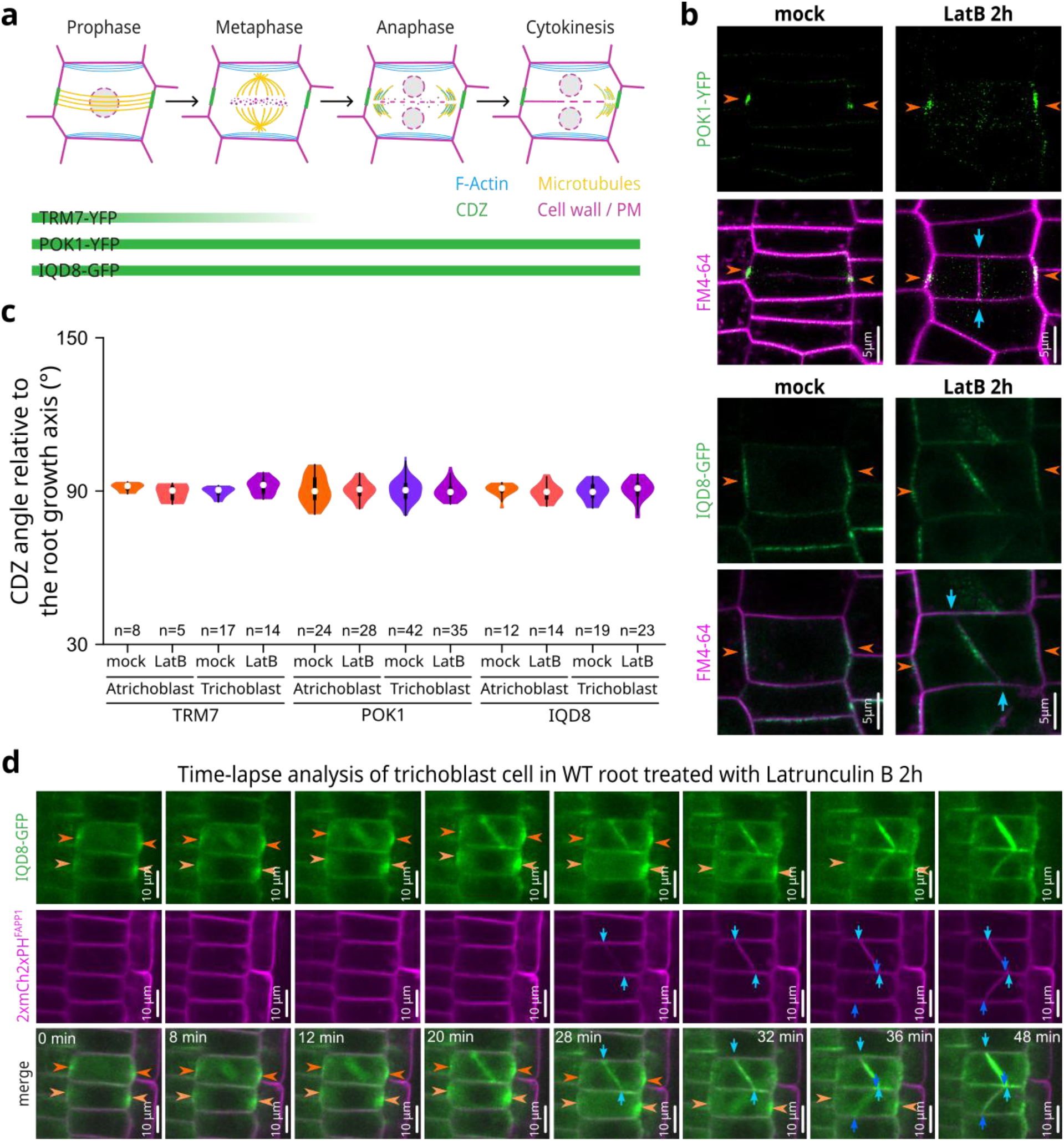
F-actin perturbation disrupts alignment between the cortical division zone and the site of cell division. **a.** Schematic representation of the spatiotemporal localisation patterns of TRM7, POK1 and IQD8 at the CDZ (green) and cell plate fusion site across different step of the cytokinesis, as indicated by green bars. **b.** Representative images of dividing trichoblasts showing the localisation of the CDZ markers POK1 and IQD8 (green) after 2h-LatB in roots stained with FM4-64 (magenta). **c.** Angles between the CDZ and root longitudinal axis after 2h-mock or -LatB. **d.** Time-lapse analysis of dividing trichoblast in root expressing *IQD8-GFP* (green) and *2xmCh2xPH^FAPP1^* (magenta), after 2h-LatB. Orange arrowheads indicate CDZ position, light blue ones the site of the cell plate attachment.

Time-lapse imaging of Arabidopsis root meristem cells expressing *IQD8-GFP* and a plasma membrane marker *2mCH-2xPH*^*FAPP1*36^showed that cells subjected to shortterm LatB treatment frequently divided along abnormal, apicobasal planes (Fig. 4d, Videos S3 and S4). However, throughout the imaging period, IQD8-GFP remained localized to the lateral CDZ, even as the cell plate formed in a perpendicular orientation. These results, consistent with previous findings, demonstrate that disruption of the actin cytoskeleton uncouples the spatial cues for phragmoplast guidance from CDZ positioning^16^—suggesting that actin is essential not for CDZ specification, but for its functional translation into accurate division execution.

### F-Actin maintains spindle orientation againt cellular geometry

To determine when actin disruption affects division orientation, we examined mitotic figures in WT 2h-mock, -LatB, and *act7-4* trichoblasts. Microtubules, the nuclear envelope, and cell walls were visualized using immunostaining and calcofluor labelling (Fig. 5a). In mock-treated WT, spindle poles aligned with the lateral PPB, consistent with proper division orientation. In contrast, LatB-treated and *act7-4* cells showed clear spindle misalignment by metaphase, with the spindle axis rotated relative to the PPB/CDZ. By integrating data from both fixed and live-imaged cells, we found that spindle poles consistently remained closely associated with the apicobasal cell faces of the dividing cell (Fig. 5b and Data S1g). These results indicate that actin integrity is required to maintain spindle orientation in case of geometrical constrains.

**Fig. 5.**
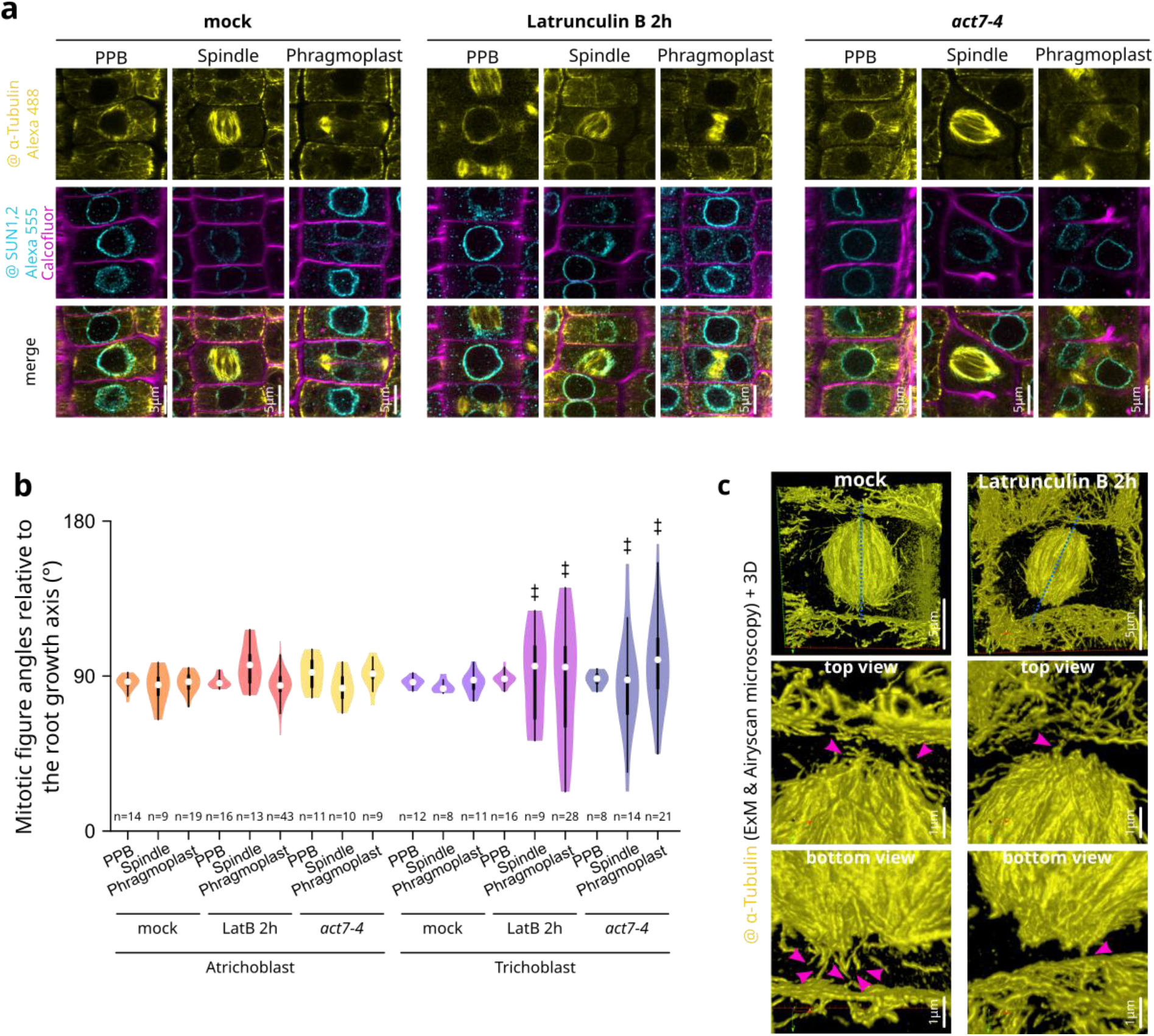
F-Actin is essential in maintaining the orientation of the spindle during plant cell division. **a.** Representative images of the mitotic figures after immunolabelling of the microtubules (@–α–tubulin, yellow), the nuclear envelope (@SUN1,2 cyan) and the cell wall staining with calcofluor (magenta) in WT mock, WT LatB-2h and *act7-4* trichobast. **b.** Angles between the microtubule’s structures and the main longitudinal axis of the root in WT 2h-mock, -LatB and *act7-4* trichobast. *N*=3. **c.** Representative 3D reconstructions of expansion microscopy (ExM) images of the spindle after the immunolabelling of the microtubules (@–α–tubulin, yellow) in in WT 2h-mock, -LatB trichobast. The top and bottom parts of the spindle pole are magnified. Pink arrowheads highlight physical connections between the microtubules growing out from the spindle poles and the apicobasal cell’s cortex. Scale bars refer to the original biological scale (expansion factor is approximately 3.7; physical size post-expansion = 18.5 μm). *N*=4. ‡ significant differences in variance distribution.

Immunolabeling of spindles microtubules (anti–α–tubulin) coupled with expansion microscopy and Airyscan imaging revealed discrete microtubule connections between the spindle poles and the apicobasal cell’s cortex in WT mock-treated cells (Fig. 5c and Video S5). These connections were markedly reduced or absent upon F-actin disruption (2h-LatB), coinciding with a spindle orientation tilt (Fig. 5c and Video S6). These findings suggest that F-actin may function to anchor the microtubule cytoskeleton to the cell cortex following PPB disassembly.

Together our data suggest that peripheral F-actin, particularly at the apicobasal domains of dividing cells, serves as a cortical landmark essential for maintaining spindle orientation. Probably via spindle fibres anchoring to the cell periphery, actin ensures the fidelity of division plane positioning against cell geometry.

## Conclusion

In multicellular organisms, morphogenesis arises from a dynamic interplay of processes, including cell shape changes, intercalation, delamination, programmed cell death, and cell division. However, in plants, particularly in the young tissues, morphogenesis is primarily driven by differential growth, as cell migration and programmed cell death are largely absent ^37^. As a result, the spatial orientation of cell division plays a pivotal role in maintaining tissue architecture and shaping organs.

Here, we identify a central role for the actin cytoskeleton in controlling division orientation in *Arabidopsis* root epidermis. Actin dynamics enable cells to integrate additional positional information and sometimes override default geometric division cues. Disruption of actin leads to misalignment between the PPB and the actual division site, revealing a disconnect between early cortical markers and final division outcomes. We further show that peripheral F-actin stabilises spindle orientation, particularly at apicobasal domains, likely by anchoring spindle fibres at the cell cortex.

These findings uncover a previously underappreciated role for actin in guiding plant cell division and suggest that cytoskeletal regulation enables robust morphogenesis by adapting division orientation to local developmental and geometrical context.

## Methods

### Growth conditions and plant materials

This study used Arabidopsis thaliana Columbia-0 (Col-0) accession as a wild-type (WT) reference genomic background. The list and description of the mutants and reporter lines used in this study can be found in Table S1. Plants were grown in soil under long-day conditions at 21 °C, 70% humidity, and 16 hours of daylight. A vapor-phase sterilization method was used for the surface sterilization of the seeds. Seeds were sown in vitro on half Murashige and Skoog (½ MS) basal medium supplemented with 0.7% plant agar and 1% sucrose (pH 5.7), grown vertically in continuous light conditions at 21 °C for six days before imaging.

### Pharmacological treatments

The perturbations of the actin cytoskeleton were studied using Latrunculin B or Cytochalasin D. Plants were vertically grown for five days in ½ MS supplemented with 0.7% plant agar and 1% sucrose (pH 5.7) and then transferred for 2 or 16 hours to growth me-dia supplemented with either 1μM Latrunculin B (Sigma) or 5μM Cytochalasin D (Sigma).

### Live cell imaging

Automated imaging of cell division in growing Arabidopsis roots expressing fluorescent proteins was primarily performed as described^38^. Six-day-old Arabidopsis seedlings were transferred to an observation chambered cover glass (Lab-Tek II, www.thermoscientific.com) containing ½ MS-0.8% agar. The plants were left to grow in their new environment in a growth chamber for two hours, with an inclination of ± 70° angle, so seedlings could acclimate to the observation chamber and keep growing. The sample was consequently placed under the spinning disk confocal microscope, and cells in the meristematic region of the root tip were subjected to time-lapse imaging. Up to three roots were observed simultaneously, and images were collected at different Z-positions every three or four minutes for up to 6 hours. The spinning disk confocal micro-scope setup was composed of an inverted Zeiss microscope (AxioObserver Z1, Carl Zeiss Group, http://www.zeiss.com/) equipped with a spinning disk module (CSUW1-T3, Yokogawa, www.yokogawa.com) and a Camera Prime 95B (www.photometrics.com) using a 63x Plan-Apochromat objective (numerical aperture 1.4, oil immersion) or an Objective LD C-Apochromat 40x/1.1 W Corr M27. GFP was excited with a 488 nm laser (150 mW), and fluorescence emission was filtered by a 525/50 nm BrightLine, a single-band bandpass filter (Semrock, http://www.semrock.com/). RFP and TdTomato were excited with a 561-nm laser (80 mW), and fluorescence emission was filtered by a 609/54-nm BrightLine single-band band-pass filter (Sem-rock). Care was taken to use similar confocal settings when comparing fluorescence intensity or for quantification.

For the analysis of actin (LifeAct-YFPv), microtubules (GFP-MBD) and CDZ (TRM7-YFPv, IQD8-GFP or POK1-YFP) markers, to visualize cell contour, seedlings were incubated in wells containing 1μM FM4-64 (Life Technologies T-3166; from a stock solution of 1.645 mM = 1 mg/ml in DMSO) in ½ MS liquid basal medium for 1 minute in the dark. Seedlings were then mounted in the same medium. High-resolution imaging was performed using a LSM980 Airyscan2 (ZEISS) using a 40× objective (numerical aperture 1.3, oil immersion). Images were acquired by sequential line switching, allowing the separation of channels by both excitation and emission. GFP, YFPv and FM4-64 were excited using 488-, 514-and 561-nm lasers, respectively. All the signals were recorded with an airyscan 2. Airyscan processing was performed with ZEN imaging software by default settings (ZEN blue).

### Calcofluor staining and immunolocalization imaging

The root meristem cell walls were stained using the calcofluor dye, following the protocol described in^39^. Seedlings were incubated overnight in a fixation buffer (50% methanol, 10% acetic acid, and 40% distilled water). The seedlings were rehydrated in ethanol baths for 10 min each: 50% ethanol, 30% ethanol, and distilled water twice. Afterward, the seedlings were transfered to the staining solution for overnight incubation (90% of clearsee solution (5% of urea, 15% of deoxycholic acid, and 10% of xylitol in distilled water) and 10% of calcofluor white solution (500 mg Fluorescent Brightener 28 in distilled water (qsp 50mL) and one drop of NaOH 10N)). Before imaging, the seedlings were rinsed for 15 minutes in clearsee solution baths. Imaging was performed on an LSM980 Airyscan2 (ZEISS) confocal microscope using a 40x objective (numerical aperture 1.3, oil immersion). Fluorescent Brightener 28 (Calcofluor) was excited using a 405nm la-ser, and whole root z-stacks were performed with 0.8 μm space between acquisitions.

For immuno-localization, seedlings were fixed in 4% paraformaldehyde and 0.1% Triton X-100 in ½ MTSB buffer [25 mM Pipes, 2.5 mM MgSO4, and 2.5 mM EGTA (pH 6.9)] for 1 hour under vacuum and then rinsed in phosphate-buffered saline (PBS) 1X for 10 min. Samples were then permeabilized in ethanol for 10 min and rehydrated in PBS for 10 min. Cell walls were digested using the following buffer for 1 hour: 2 mM MES (pH 5), 0.20% driselase, and 0.15% macerozyme. Tissues were incubated overnight at room temperature with the B-5-1-2 monoclonal anti-α-tubulin (Sigma-Aldrich) and the anti-SUN1,2 (Agrisera) antibody. The next day, tissues were washed for 15 min in PBS and 50 mM glycine, incubated with secondary antibodies (Alexa Fluor 555 goat anti-rabbit for SUN1,2 antibody and Alexa Fluor 488 goat anti-mouse for the tubulin antibody) overnight, and washed again in PBS and 50 mM glycine. Samples were incubated in 10% calcofluor white solution for 2 hours and then mounted in VECTASHIELD.

High-resolution imaging was performed using a LSM980 Airyscan2 (ZEISS) using a 40× objective (numerical aperture 1.3, oil immersion). Multi-color images were acquired by sequential line switching, allowing the separation of channels by both excitation and emission. Alexa Fluor 555, Alexa 488 and Fluorescent Brightener 28 (Calcofluor) were excited using 561-, 488-, and 405-nm lasers, respectively. All the signals were recorded with an airyscan 2. Airyscan processing was performed with ZEN imaging software by default settings (ZEN blue).

### Growth conditions and plant materials

This study used Arabidopsis thaliana Columbia-0 (Col-0) accession as a wild-type (WT) reference genomic background. The list and description of the mutants and reporter lines used in this study can

### Expansion microscopy

The protocol used was recently described^40^. Briefly, seedlings were fixed in 4% paraformaldehyde dissolved in MTSB buffer [50 mM PIPES, 5 mM EGTA, 5 mM MgSO4 pH 7 with KOH] for 1 hour and washed with MTSB buffer. The root tips were cut and transferred onto a Corning BioCoat Poly-D-lysine glass coverslip 12 mm (VWR) and allowed to dry for approximately 1 hour. The immunolabeling procedure is the same as described in ^41^, except for the cell wall digestion incubation time that was set up to 40 min to optimize expansion. The following antibodies were used: mouse anti-α-tubulin (T5168, Merck, dilution 1/1000), goat anti-mouse-Alexa Fluor 594 (A-11005, Invitrogen, dilution 1/500).

After immunolabeling, the samples were incubated in the anchoring solution [0.1 mg/mL Acryloyl-X-SE (Thermo Fisher Scientific) in 1× PBS buffer] overnight at RT, washed with 1× PBS buffer, and incubated in the monomer solution [11.7% NaCl, 3% acrylamide, 0.15% N,N′-methylenebisacrylamide, 8.625% sodium acrylate, and 0.5% 4-hydroxy-tempo] overnight at 4 °C. Samples were then preincubated in the gelling solution [0.2% ammonium persulfate and 0.2% TEMED in 1 mL of monomer solution] for 30 min on ice under gentle agitation. Coverslips were transferred upside down onto a drop of 35 mL of the freshly prepared gelling solution in a humid chamber with parafilm and incubated for 2 hours at 37 °C. After gelation, the embedded samples were incubated in protein digesting solution [50 mM Tris (pH 8.0), 2.5 mM EDTA, 0.5% Triton X-100, 0.8M Guanidine-HCl, and 8 units/mL Proteinase K] at RT overnight. The excess of gel around the sample was removed using a razorblade. Gels were then incubated for 2 hours at 30 °C in 2% driselase solution in 1× PBS buffer for an additional cell wall digestion. Gels were transferred into a 60 mm round petri-dish and expanded in excess of ddH20. The ddH2O was renewed every 45 min until the plateau was reached (around 3 hours). Imaging was performed using an LSM880 (ZEISS) with a 40× objective (numerical aperture 1.2, water diping). Alexa Fluor 594 was excited using a 594-nm laser, and optical sections were collected with 0.38 μm space between acquisitions. The signals were recorded using an Airyscan detector in super-resolution mode. Airyscan processing was performed with ZEN imaging software (ZEN blue). 3D reconstruction was performed using Icy software.

### Dividing cell segmentations and simulations

The pipeline used for cell division segmentation and predictions used in this study was described previously^21^. For every condition/genotype, ten recently divided trichoblast and atrichoblast root cells were selected from calcofluor staining images and 3D segmented in MorphographX software^23^. Mother cells were reconstructed by combining the two daughter cells. 500 cell divisions in reconstructed mother cells were simulated using the model introduced by ^25,26^. This model randomly draws the volume ratio (VR: volume smaller daughter cell / reconstructed mother cell) between 0.2 and 0.5 and generates a partitioning of the mother cell that locally minimizes the interface area between the two daughter cells. For analysis, only symmetrical simulated divisions with VR values within the empirically observed range (0.42 < VR < 0.50), representing ∽30% of the total simulations per cell, were considered. The relative interface area index was calculated as follow:

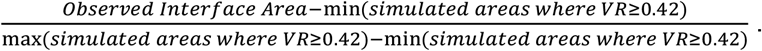

### Measures, counting, and statistical analysis

For quantitative imaging, pictures of epidermal root meristem cells were taken with detector settings optimized for low background and no pixel saturation. Care was taken to use similar confocal settings when comparing fluorescence intensity.

Cell division angles were quantify as previously described in ^21^ using custom-made FIJI macro (MacroDivRootGascon_et_al2024V1.ijm) available at https://github.com/RDP-vbayle/SiCE_FIJI_Macro/tree/main/misc. To assess the effect of pharmacological treatments on the actin polarization index, measurements were made using ImageJ according ^42^. The cortical division zone (CDZ) and microtubules mitotic figures’ angles were manually measured using ImageJ, “straight line”, and “angle tool”.

Statistical analysis where performed in R (v. 4.4.0; R Core Team, 2019), using R studio interface and the packages ggplot2 ^43^. Graphs were obtained with R and R-studio software and customized with Inkscape (https://inkscape.org). In the supplementary data related to every figure are mentioned the specific test performed to do the comparisons.

## Acknowledgements

We are grateful to the Signalisation and Endomembrane SiCE group (RDP, ENS de Lyon, France) for their comments and discussions. We are grateful to Dr Charlotte Kirchhelle (RDP, ENS de Lyon, France) for her insightful feedback on the manuscript. We thank Patrice Bolland and Alexis Lacroix from our plant facility (RDP, ENS de Lyon, France), and Claire Lionnet (RDP, ENS de Lyon, France) for the advice on microscopy. We acknowledge the contribution of SFR Biosciences (UMS3444/CNRS, US8/Inserm, ENS de Lyon, UCBL) facilities at the LBI-PLATIM-MICROSCOPY for assistance with imaging. We thank Laure Mancini (RDP, ENS de Lyon, France) for the help running cell division predictions and J. We are grateful to Johanna Dickmann (RDP, ENS de Lyon, France) for the double line LifeAct-TdTomato Lti6b-GFP. We thank Dr. Abidur Rahman (Department of Plant Biosciences, Faculty of Agriculture, Iwate University, Morioka, Japan) for sharing the actin mutants (*act2-1, act7-4, act7-6, act8-2, act2-1act8-2* and *act7-4act8-2*).

## Fundings

National Agency for Research, ANR PlantScape, ANR-20-CE13-0026-02 (DB, PA, MCC). National Agency for Research, (ANR-DFG DIVFUSE, ANR-22-CE92-0038-02 (YB, MCC). IJPB benefits from the support of Saclay Plant Sciences-SPS (ANR-17-EUR-0007). EMBO fellowship, ATLF 646-2021 (European Molecular Biology Organization, Heidelberg, DE) (CG).

### Author contributions

Conceptualization: MCC, CG; Methodology: CG, MP, SM, YS, GC, VB, MG, YJ, DB, YB, MP, PA and MCC; Investigation: CG, MP, SM, YS, GC, VB, MG; Funding acquisition: CG, YJ, DB, YB, MP, PA and MCC; Supervision: YB and MCC; Writing – original draft: MCC, CG; Writing – review & editing: CG, MP, YJ, VB, YB, MP, PA and MCC.

## Materials & Correspondence

All data are available in the manuscript or the supplementary materials and are available upon request.

### Supplementary figures

**Fig. S1.**
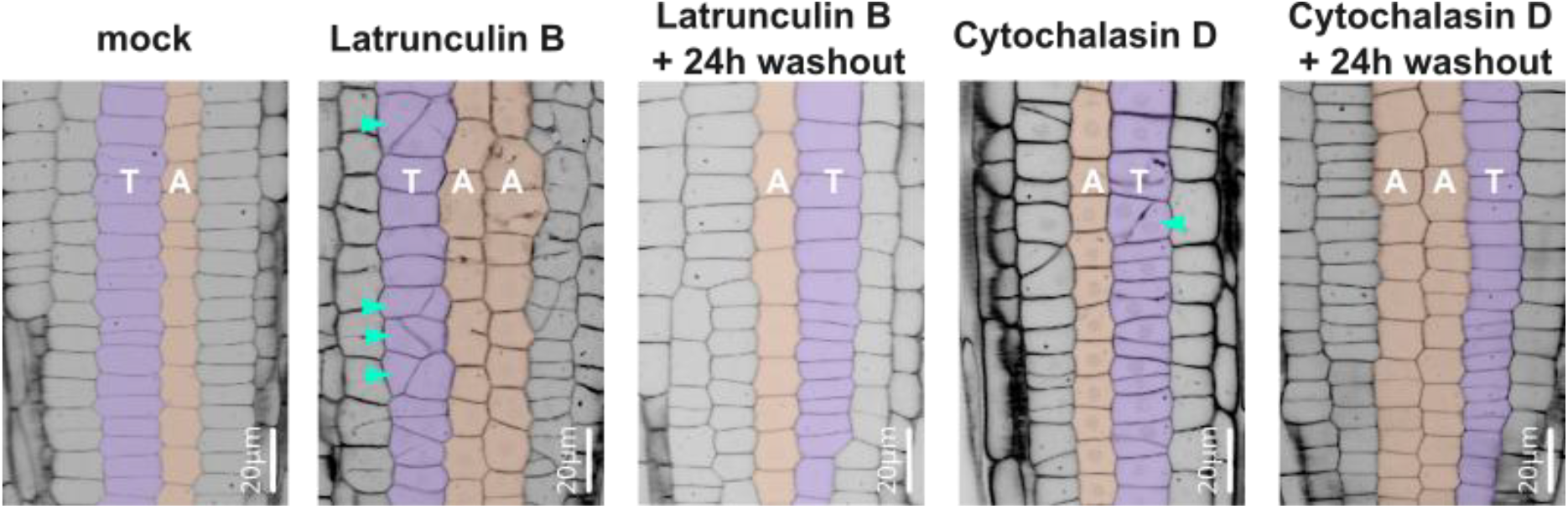
A representative image of the epidermis cell layer view was extracted from a Z-stack of calcofluor-stained fixed wild-type roots (6-day-old) treated for 16h with DMSO (mock), 1μM Latrunculin B or 5μM Cytochalasin D. Atrichoblast (A) cell layers are indicated in orange, while trichoblast (T) cells are in purple. Turquoise arrowheads point abnormal apico-basal divisions.

**Fig. S2.**
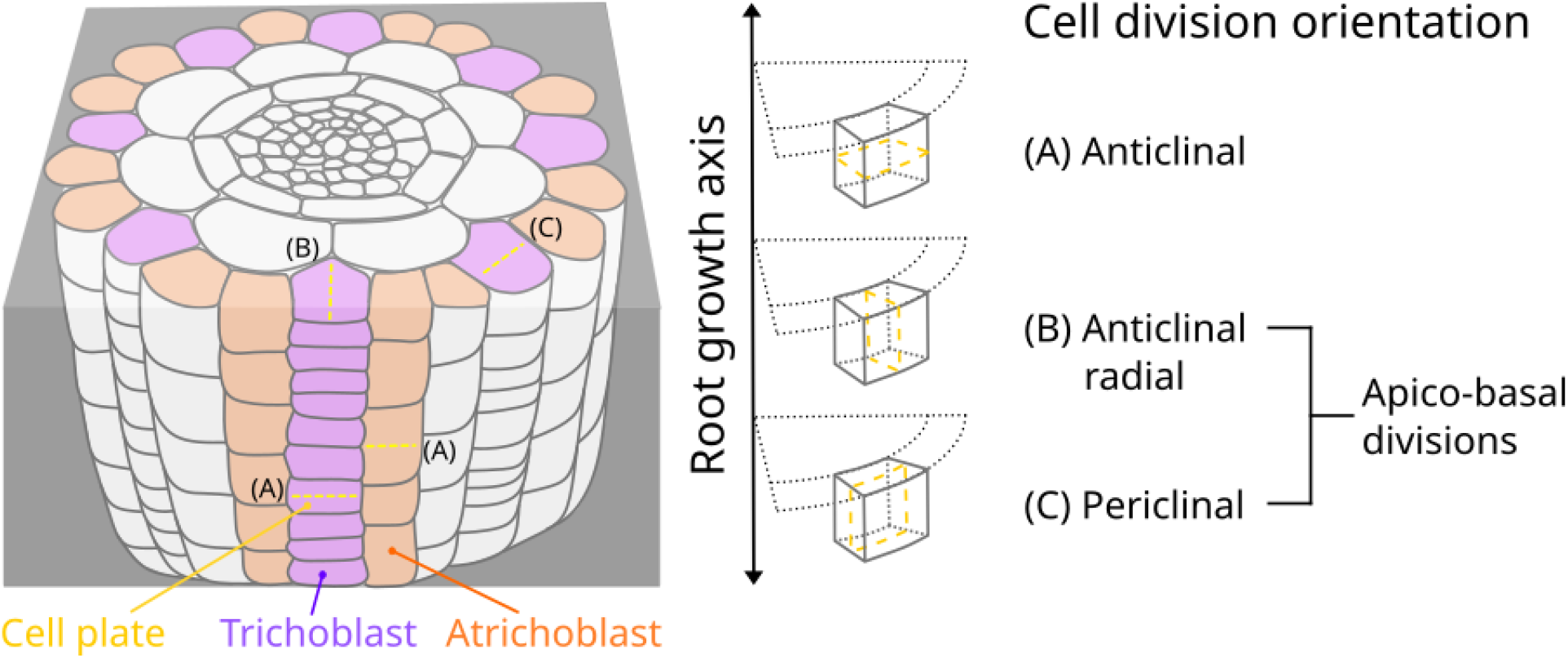
A 3D schematic representation of a root depicting the possible cell division orientations and showing those classified as apico-basal.

**Fig. S3.**
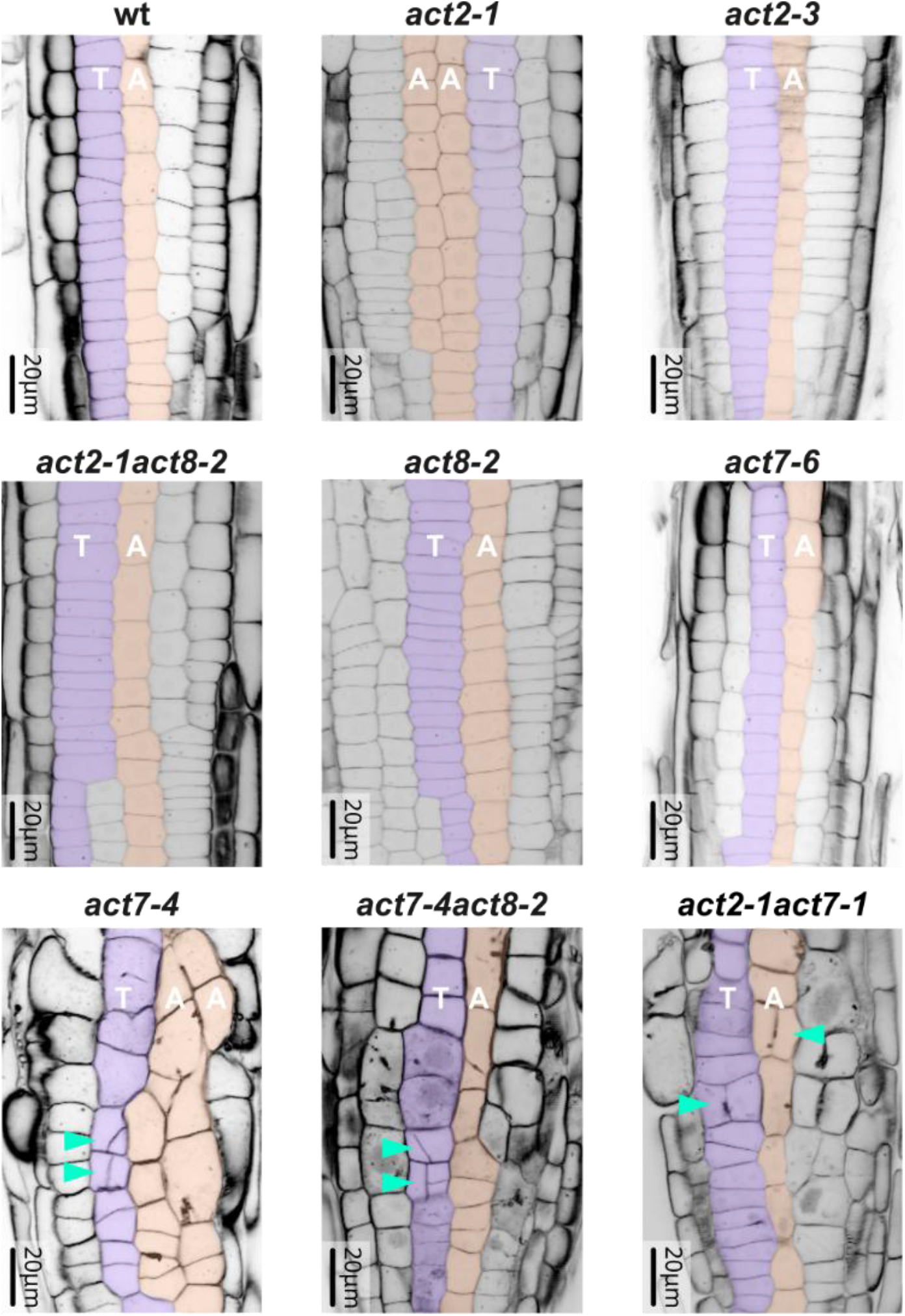
A representative image of the epidermis cell layer view was extracted from a Z-stack of calcofluor-stained fixed roots (6-day-old) of actin mutants. Atrichoblast (A) cell layers are indicated in orange, while trichoblast (T) cells are in purple. Turquoise arrowheads point abnormal apico-basal divisions.

**Table S1.** List of plant materials used.

| Line | Identifier | Source |
| --- | --- | --- |
| Col-0 (WT) | EW 20-640 | ABRC |
| *trm678* | N/A | 7 |
| *act7-4* | N/A | 32,32,33 |
| *act2-1* | N/A | 32,32,33 |
| *act8-2* | GABi-Kat line 480C07 | 33,44 |
| *act2-1 act8-2* | N/A | 33,44 |
| *act7-4,act8-2* | N/A | 33,44 |
| *act7-6* | SALK_131610C | 33,45 |
| *act2-1,act7-1* (Ws ecotype) |  | 32 |
| Lifeact-YFPv |  | 46 |
| *pPOK1:YFP-POK1* |  | 47 |
| GFP-MBD | NASC N799990 | ABRC |
| RFP-MBD |  | 47 |
| mCit-P4M |  | 48 |
| 2xmCH-2xPH<sup>FAPP1</sup> |  | 36 |
| *pUb10:IQD8-GFP* |  | 35 |
| GFP-MBD x 2xmCH-2xPH<sup>FAPP1</sup> |  | 21 |
| RFP-MBD x mCit-P4M |  | 21 |
| IQD8-GFP x 2xmCH-2xPH<sup>FAPP1</sup> |  | this study |
| RPS5apro:Lifeact-TdTomato |  | 49 |
| 35S:Lti6B-GFP |  | 50 |
| TRM7pro:TRM7-3YPet |  | 7 |
| 35S:GFP-Lti6B x LifeAct-tdTOM |  | this study |

**Video S1.**

Time-lapse analysis of actin apico basal polarization in an atrichoblast cell co-expressing *LifeActtdTom* (cyan) and *Lti6b-GFP* (magenta) markers in 6-day-old wild type plants. Movie corresponding to Fig. 2a.

**Video S2.**

Time-lapse analysis of actin apico basal polarization in a trichoblast cell co-expressing *LifeActtdTom* (cyan) and *Lti6b-GFP* (magenta) markers in 6-day-old wild type plants. Movie corresponding to Fig. 2a.

**Video S3.**

Time-lapse analysis of a WT root co-expressing the CDZ marker *IQD8-GFP* (green) and *2xmCH2xPH^FAPP1^*(magenta) in 6-day-old plants. Movie corresponding to Fig. 4d.

**Video S4.**

Time-lapse analysis of a WT root treated 2h with Latrunculin B co-expressing the CDZ marker *IQD8-GFP* (green) and *2xmCH-2xPH^FAPP1^*(magenta) in 6-day-old plants. Movie corresponding to Fig. 4d.

**Video S5.**

Representative 3D reconstructions of expansion microscopy (ExM) images of the spindle after the immunolabelling of the microtubules (using @–α–tubulin, yellow) in trichoblast cell from WT mock treated plant (6-day-old). Movie corresponding to Fig. 5c.

**Video S6.**

Representative 3D reconstructions of expansion microscopy (ExM) images of the spindle after the immunolabelling of the microtubules (using @–α–tubulin, yellow) in trichoblast cell from WT LatB-2h treated plant (6-day-old). Movie corresponding to Fig. 5c.

**Data S1. (separate file)**

Quantification and statistical analysis, related to Figures 1–5.

a. Data S1a. Relative to Fig. 1b, 1d, 1g and 1h.
b. Data S1b. Relative to Fig. 1f.
c. Data S1c. Relative to Fig. 2c.
d. Data S1d. Relative to Fig. 2e.
e. Data S1e. Relative to Fig. 3a and 3c.
f. Data S1f. Relative to Fig. 4c.
g. Data S1g. Relative to Fig. 5b.

**Data S2. (separate file)**

Cell division predictions, related to Fig. 1c and 1h. WT atrichoblast cell 1-10, WT trichoblast cell 1-10, *trm678* atrichoblast cell 1-10 and *trm678* trichoblast cell 1-10 (6-day-old plants).

**Data S3. (separate file)**

Cell division predictions, related to Fig. 3b. Atrichoblast cell 1-10 and trichoblast cell 1-10 from WT 16h-DMSO treated (mock), WT 16h-LatB treated, *act7-4* and *trm678* 16h-LatB treated plants (6-day-old).

